# An explainable artificial intelligence framework reveals mutations associated with drug resistance in *Mycobacterium tuberculosis*

**DOI:** 10.1101/2024.12.18.629160

**Authors:** Hui Cen, Peng Zhang, Yunchao Ling, Guoping Zhao, Guoqing Zhang

## Abstract

Understanding the mechanisms of drug resistance in *Mycobacterium tuberculosis* (MTB) is essential for the rapid detection of resistance and for guiding effective treatment, ultimately contributing to reducing the global burden of tuberculosis (TB). Under anti-TB drugs pressure, MTB continues to accumulate resistance loci. The current repertoire of known resistance-associated mutations requires further refinement, necessitating efficient methods for the timely identification of potential resistance sites. Here, we introduce xAI-MTBDR, an explainable artificial intelligence framework designed to identify new resistance-associated mutations and predict drug resistance in MTB. It outperforms state-of-the-art methods in predicting drug resistance for all first-line drugs, and scoring each mutation’s contribution to resistance. By leveraging public whole-genome sequencing data from nearly 40,000 MTB isolates, the framework identified 788 candidate resistance-related mutations and revealed 30 potential resistance markers, several of which are structurally located closer to their respective drugs compared to known resistance mutations. Furthermore, these scores enabled the framework to efficiently subgroup isolates with different resistance mechanisms and reflect varying levels of resistance. The framework serves as a valuable tool for accurate detection of drug-resistant MTB and offers new insights into its underlying mechanisms.

## Introduction

Tuberculosis (TB), caused by *Mycobacterium tuberculosis* (MTB), is the world’s leading bacterial infectious disease^1,2^. In 2022, approximately 10.6 million people developed TB, resulting in 1.3 million deaths^3^. The rise of drug-resistant TB has worsened this crisis, with treatment success rates for drug-resistant TB being 25% lower than for drug-susceptible TB^3,4^. This highlights the urgent need to understand resistance mechanisms for rapid detection of drug-resistant MTB, guiding effective treatment and reducing the global TB burden^5^.

The majority of drug resistance in MTB isolates is attributed to chromosomal mutations^6^. Genome-wide association studies (GWAS) have been successfully applied to investigate novel resistance mechanisms in MTB by identifying nucleotide variants associated with drug resistance^7^. This approach has not only detected known resistance loci but also uncovered new resistance-related loci, such as specific sites in the *ald* gene correlated with increased resistance to D-cycloserine, novel codons in the *ethA* gene associated with ethionamide resistance, and variations in the promoter region of *thyX* related to para-aminosalicylic acid resistance^8-10^. The World Health Organization (WHO)-endorsed catalogue of MTB resistance mutations, based on whole-genome sequencing (WGS) data, categorizes mutations according to their association with resistance and serves as a high-quality reference^11,12^. Pre-determined mutation catalogues, which are essential for detecting drug resistance, form the foundation upon which popular prediction tools such as TB-profiler^13^ and Mykrobe^14^ infer drug resistance. With the rapid accumulation of WGS data, these catalogues need further refinement^4,7^, particularly for low-frequency mutations in natural populations^7^.

Deep learning models have enriched mutation catalogues by identifying complex patterns in genomic data^15^, as demonstrated by Green et al.’s convolutional neural network, which identified 18 sites not previously associated with resistance^16^. However, this study did not explore deeper individual-level interpretations, which are essential for understanding how specific mutations contribute to resistance in individual cases. The need for models that provide both global (population-level) and local (individual-level) explanations is critical. Global explanations help identify broad mutation contributions, while local explanations are essential for understanding individual mutations that drive resistance in specific isolates. The SHapley Additive exPlanations (SHAP) method is well-suited for this purpose, offering both global and local insights by evaluating the contribution of each mutation to drug resistance predictions^17^. SHAP’s dual capability enhances mutation catalogue completeness and improves the understanding of resistance mechanisms, making it a valuable tool for research on pathogenic microorganisms.

In this study, we present xAI-MTBDR, an explainable artificial intelligence framework that integrates machine learning with SHAP to identify new resistance-associated mutations and predict drug resistance in MTB. Leveraging public whole-genome sequencing data from nearly 40,000 MTB isolates, xAI-MTBDR outperforms state-of-the-art methods in predicting resistance for all first-line anti-TB drugs. By providing both population-wide and individual-level insights, xAI-MTBDR enables revelation of potential biomarkers, grouping of drug-resistant isolates, and reflection of varying resistance levels. It is helpful for advancing the understanding of TB resistance mechanisms.

## Result

### The framework and datasets characteristics

The framework is consisted of four main stages: dataset collection, feature extraction, model construction and explanation (Fig.1, detailed in Methods). In stage 1, we compiled 39,145 high-quality MTB isolates from three datasets, each providing WGS and DST data for 11 anti-TB drugs^11,18,19^. In stage 2, features were derived from diverse mutations, including insertions, deletions, and single nucleotide polymorphisms (SNPs), based on 49 preselected antimicrobial resistance (AMR) genes and promoter regions^20^. Stage 3 involved training and testing for each drug by the leave-one-out (LOO) strategy. In the final stage, we applied SHAP to provide interpretable insights, identifying key mutations driving resistance and exploring resistance mechanisms at both population and individual levels.

**Fig.1.**
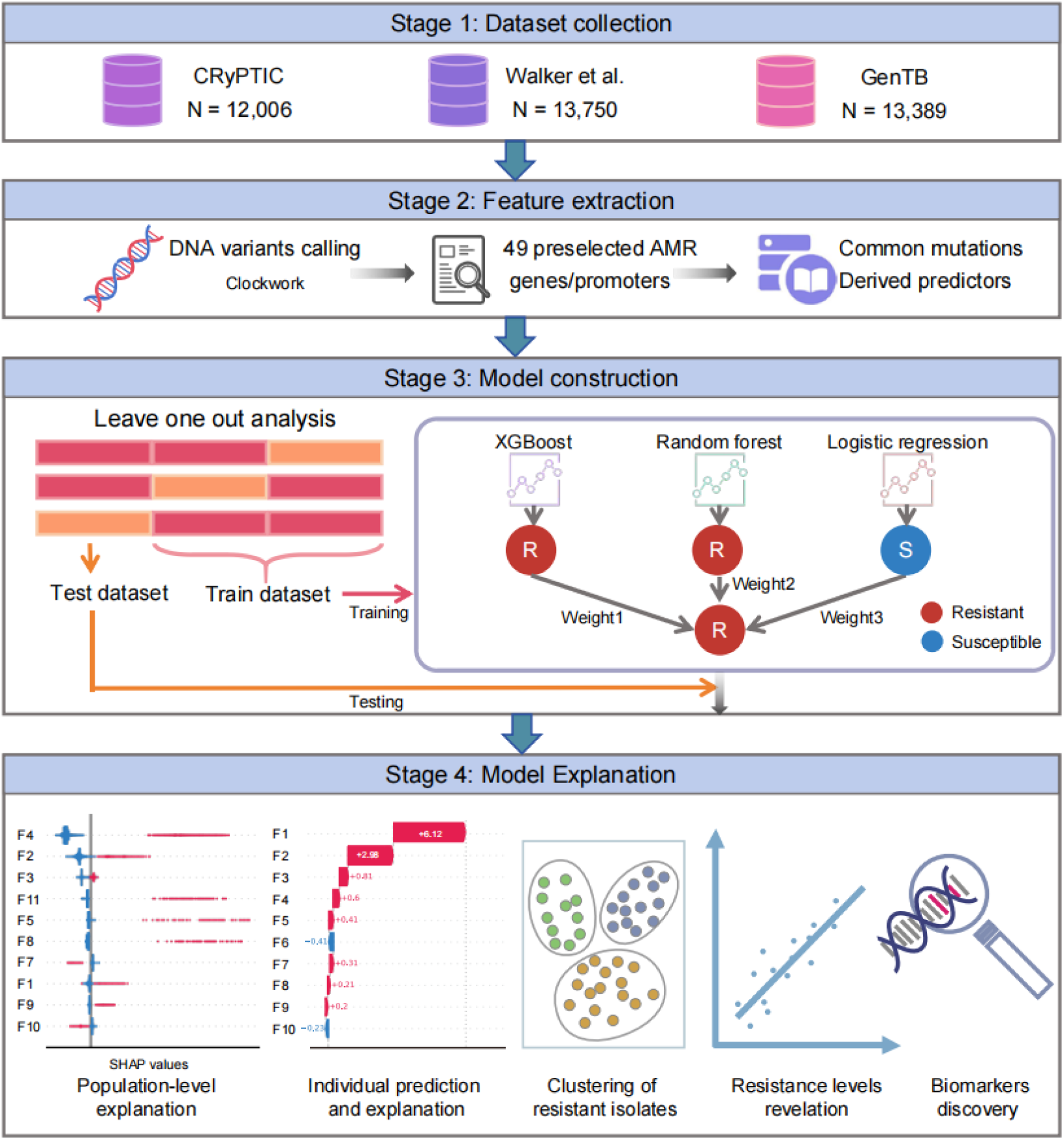
Workflow of the xAI-MTBDR framework. The process includes four stages: (1) Data collection from three datasets; (2) Variation calling and feature extraction from genes and promoter regions; (3) Construction and testing of machine learning models; and (4) Application of SHAP for model explanation.

Proportions of drug-resistant isolates across different drugs are summarized in Table 1. Each drug is represented by at least 250 isolates per dataset^16^. Rifampicin and isoniazid have the highest numbers, with 36,552 and 36,240 isolates. In contrast, capreomycin has the fewest DST entries, with only 11,246 isolates. The proportion of resistant isolates also varies, ranging from 8% for amikacin to 34% for isoniazid.

**Table 1.** Proportions of drug-resistant isolates among all isolates.

| Drugs | Datasets | CRyPTIC | Walker et al. | GenTB | All |
| --- | --- | --- | --- | --- | --- |
| 1<sup>st</sup> line drugs |  |  |  |  |  |
| RIF |  | 39% (4564/11672) | 18% (2120/11732) | 22% (2832/13148) | 26% (9516/36552) |
| INH |  | 48% (5589/11524) | 26% (3137/11876) | 27% (3494/12840) | 34% (12220/36240) |
| EMB |  | 25% (2703/10737) | 9% (961/10584) | 15% (1554/10596) | 16% (5218/31917) |
| PZA |  | 32% (309/961) | 13% (725/5410) | 12% (1319/10677) | 14% (2353/17048) |
| 2<sup>nd</sup> line drugs |  |  |  |  |  |
| AMK |  | 7% (859/11751) | 7% (267/3997) | 14% (504/3705) | 8% (1630/19453) |
| CAP |  | 9% (272/3195) | 6% (251/4260) | 13% (507/3791) | 9% (1030/11246) |
| ETH |  | 18% (1960/10970) | 29% (915/3122) | 39% (245/623) | 21% (3120/14715) |
| KAN |  | 9% (1079/11745) | 8% (358/4252) | 15% (452/2919) | 10% (1889/18916) |
| LEVO |  | 17% (2014/11648) | 19% (1016/5294) | 18% (418/2381) | 18% (3448/19323) |
| MXF |  | 12% (1360/11538) | 17% (749/4282) | 13% (291/2170) | 13% (2400/17990) |
| STR |  | 46% (1066/2329) | 24% (1279/5253) | 30% (1701/5617) | 31% (4046/13199) |

RIF, rifampicin; INH, isoniazid; EMB, ethambutol; PZA, pyrazinamide; AMK, amikacin; CAP, capreomycin; ETH, ethionamide; KAN, kanamycin; LEVO, levofloxacin; MXF, Moxifloxacin; STR, streptomycin. The WHO classifies these as first-line and second-line anti-TB drugs<sup>36</sup>.

### xAI-MTBDR outperforms state-of-the-art methods in predicting resistance

Based on the xAI-MTBDR framework, we trained individual models for each drug to predict MTB resistance and compared it with four state-of-the-art methods, including machine learning models (random forest and logistic regression), deep learning models (WDNN), and catalog-based methods (TB-profiler)^13,20,21^. As shown in Table 2, our models improved area under the curve (AUC) by 0.271-1.439% for first-line drugs, with an average improvement of 0.564%, and by 0.045-3.060% for second-line drugs, with an average improvement of 1.135%. The lower performance for amikacin and capreomycin may be due to the limited number of isolates. These results demonstrate that our framework outperforms existing methods in predicting drug resistance.

**Table 2.** The prediction performance (AUC) of methods for different drugs.

| Methods<br>Drugs | xAI-MTBDR | RF | LR | WDNN | TB-profiler | Performance |
| --- | --- | --- | --- | --- | --- | --- |
| 1<sup>st</sup> line drugs |  |  |  |  |  |  |
| RIF | **0.981** | 0.969 | 0.977 | 0.978 | 0.963 | 0.271% ↑ |
| INH | **0.957** | 0.946 | 0.955 | 0.952 | 0.908 | 0.227% ↑ |
| EMB | **0.955** | 0.946 | 0.952 | 0.943 | 0.914 | 0.318% ↑ |
| PZA | **0.916** | 0.903 | 0.899 | 0.895 | 0.684 | 1.439% ↑ |
| 2<sup>nd</sup> line drugs |  |  |  |  |  |  |
| AMK | 0.913 | **0.915** | 0.913 | 0.854 | 0.869 | - 0.311% |
| CAP | 0.902 | **0.903** | 0.891 | 0.824 | 0.827 | - 0.039% |
| ETH | **0.822** | 0.789 | 0.798 | 0.743 | 0.616 | 3.060% ↑ |
| KAN | **0.904** | 0.902 | 0.903 | 0.818 | 0.779 | 0.045% ↑ |
| LEVO | **0.941** | 0.926 | 0.936 | 0.910 | 0.919 | 0.603% ↑ |
| MXF | **0.935** | 0.919 | 0.925 | 0.904 | 0.899 | 1.056% ↑ |
| STR | **0.934** | 0.915 | 0.925 | 0.898 | 0.859 | 0.910% ↑ |

RF, random forest; LR, logistic regression; RIF, INH, EMB, PZA, AMK, CAP, ETH, KAN, LEVO, MXF, STR: see Table 1 for abbreviations.

### Population-level explanations of SHAP values aligned with WHO mutation categories

The WHO MTB resistance mutation catalogue classifies mutations into five categories based on their confidence level of association with resistance^12^. In the xAI-MTBDR framework, SHAP analysis revealed a strong correlation between mutation importance (Supplementary Fig.1a) and these categories. For rifampicin, category 1 mutations (associated with resistance) had the highest SHAP values, ranging from 4.62 to 6.29, while category 5 mutations (not associated with resistance) had the lowest (from -0.03 to 0.15) (Fig.2a). Similar trends were observed for isoniazid and ethambutol (Fig.2b,c). In category 3 (mutations with uncertain resistance associations), some mutations overlapped with category 1, particularly for isoniazid (InhA I21V, InhA S94A, KatG S315G) and ethambutol (EmbB D1024N), whose SHAP values were above the median of category 3, aligning more closely with category 1, consistent with their reported resistance associations^22-24^. These findings suggest SHAP values may help reclassify uncertain category 3 mutations (Fig.2b,c). For pyrazinamide, where the WHO mutation catalogue adopted a “relaxed” grading criteria, we observed some overlap in SHAP values across categories (Fig.2d), but category 1 mutations still showed relatively higher SHAP values. Overall, SHAP values aligned well with the WHO mutation catalogue, effectively distinguishing resistance-associated mutations at the population level.

**Fig.2.**
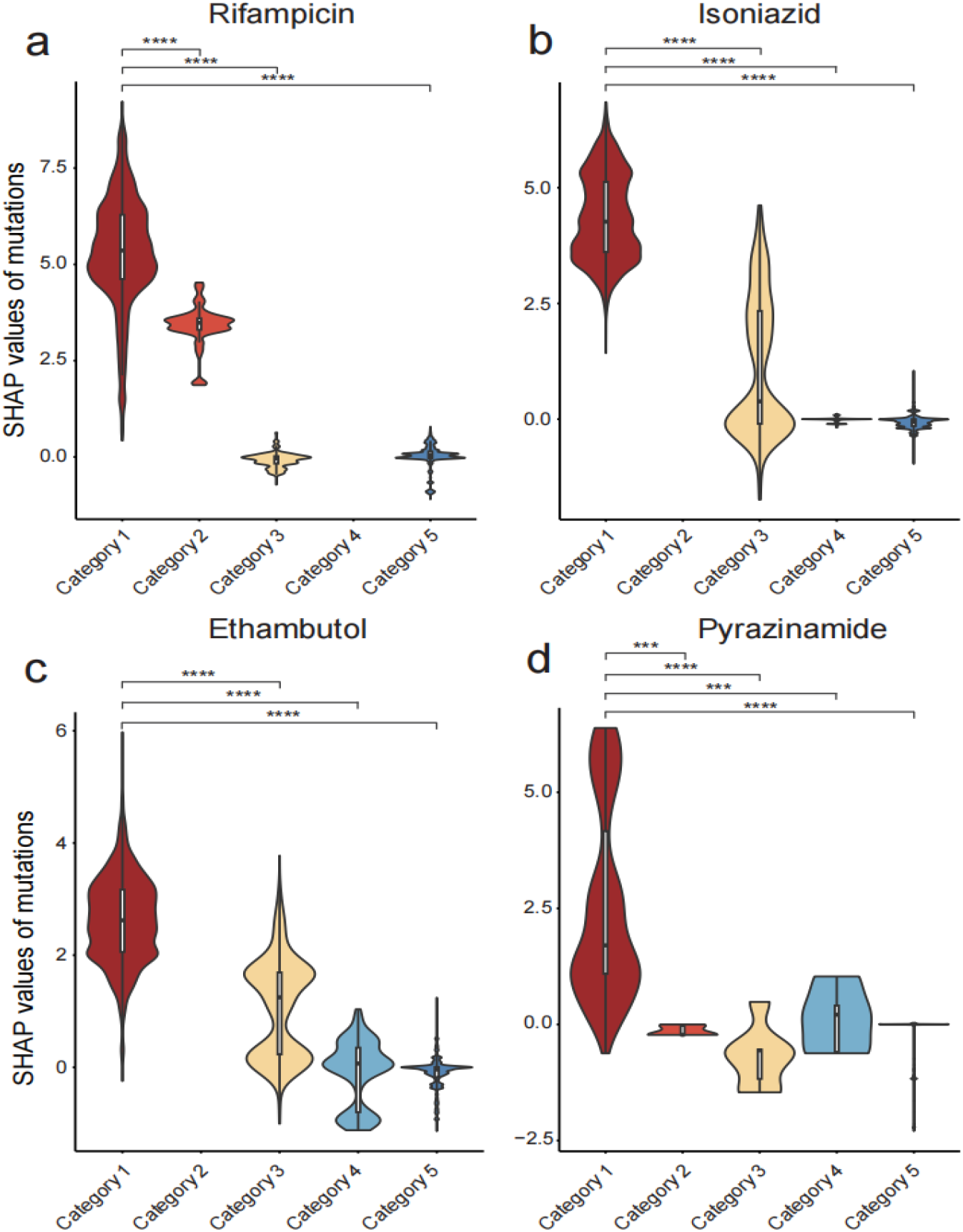
Population-level explanations for first-line drugs. **a**-**d**, Violin plot depicts the SHAP values of mutations in relation to their WHO classifications. Category 1: associated with resistance; Category 2: associated with resistance – interim; Category 3: uncertain significance; Category 4: not associated with resistance – interim; Category 5: not associated with resistance. Some mutations from the WHO mutation catalogue are not present in our model’s feature list and thus do not have corresponding SHAP values. Significance was determined using Mann-Whitney test. \*\*\**P* < 0.001; \*\*\*\**P* < 0.0001.

### Individual-level explanations identify potential resistance markers

The individual-level explanations revealed variations in mutation contributions among isolates (Supplementary Fig.1b), enabling the identification of 98 important mutations for the four first-line drugs, with 30 were not included in the WHO category 1 or 2 (Supplementary Table 1). For isoniazid and ethambutol, we identified potential resistance markers, excluding rifampicin due to the absence of novel markers in RpoB, and pyrazinamide as its markers overlapped with known mutations in PncA. The potential markers I194 (isoniazid) and D1024 (ethambutol) were located 3.7 Å and 6.1 Å from their respective drugs, compared to 4 Å and 8.6 Å for the known markers S94^25^ and M306 (Fig.3a,b, Supplementary Table 2). The median distances of known and potential markers to ethambutol were 12.6 Å and 9.1 Å, respectively, with no significant difference (Fig.3c), suggesting these mutations may influence the interaction between EmbB and ethambutol.

**Fig.3.**
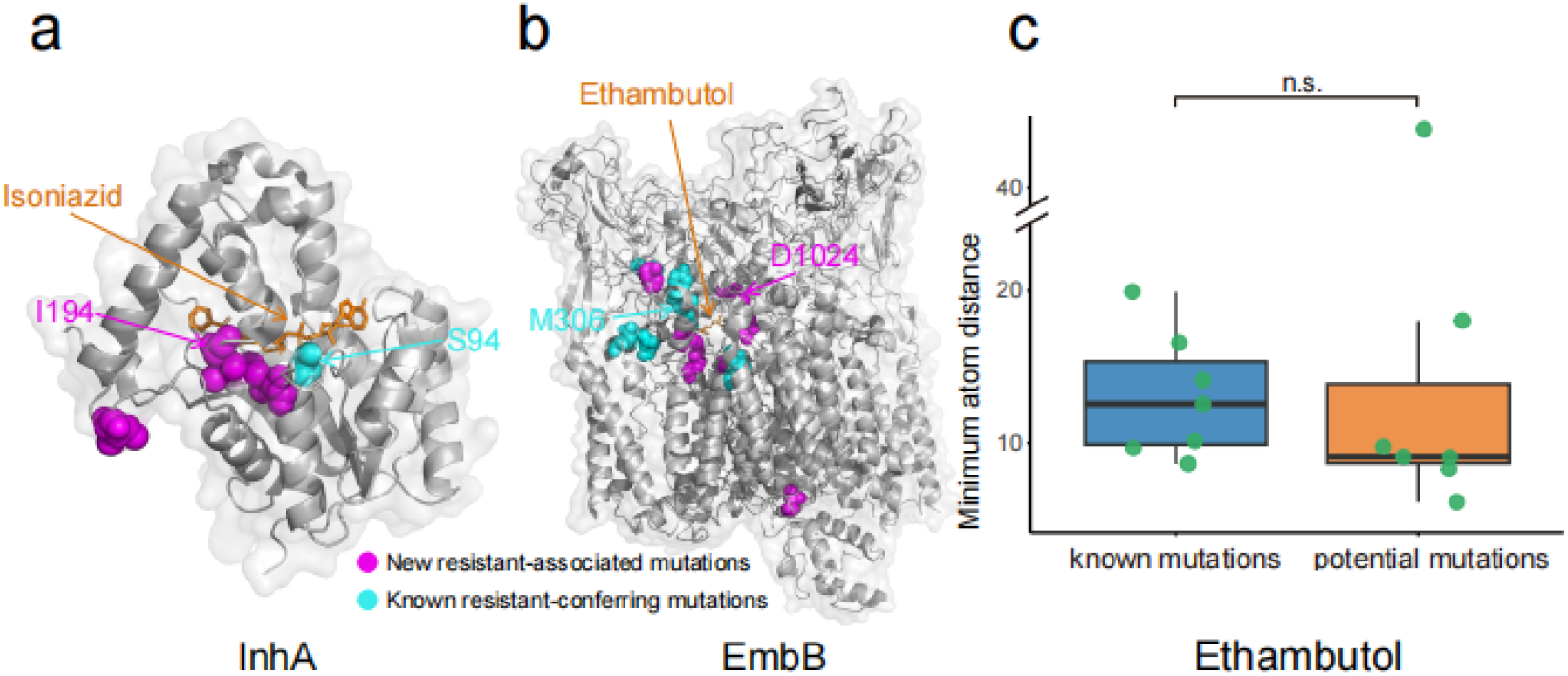
Structural analysis of resistance mutation. **a-b**, Resistance markers in the structure of InhA (PDB: 2NV6) and EmbB (PDB: 7BVF) protein. **c**, Minimum atom distance between known and potential resistance mutations to ethambutol. Mann-Whitney test was used to compare differences between the known mutations group and potential mutations group. n.s.: *P* > 0.05, not statistically significant.

### Individual-level explanations subgroup MTB drug-resistant isolates

We performed a principal component analysis (PCA) on the SHAP values of the isolates. For rifampicin, the susceptible and resistant isolates were clearly separated, and the resistant isolates were clustered into three distinct groups (Fig.4b, Supplementary Fig.2 and 3). In contrast, the PCA of the original features failed to achieve this separation (Fig.4a). Similar patterns were observed for other first-line drugs (Supplementary Fig.4). We further visualized the top 10 rifampicin resistance-associated mutations across these three groups of resistant isolates, which may reflect the underlying mechanisms driving their differentiation (Fig.4c). Group 1 is primarily driven by the RpoB D435V mutation within the 81-bp rifampicin resistance-determining region (RRDR, corresponding to codons 426–452 in MTB). Group 2, which contains the most isolates, is predominantly associated with the RpoB S450L mutation within the RRDR. In contrast, group 3 displays a more diverse mutation pattern, including non-RRDR RpoB V170F and I491F. These mutations are positioned near or directly interact with rifampicin (Fig.4d), potentially altering the structure of the active center or the interactions between RNA polymerase (RNAP) subunits, leading to resistance, consistent with previous research^26^. These results suggest that SHAP values not only capture key characteristics of drug-resistant isolates, but also reveal complex resistance mechanisms, emphasizing the importance of non-RRDR mutations in resistance detection.

**Fig. 4.**
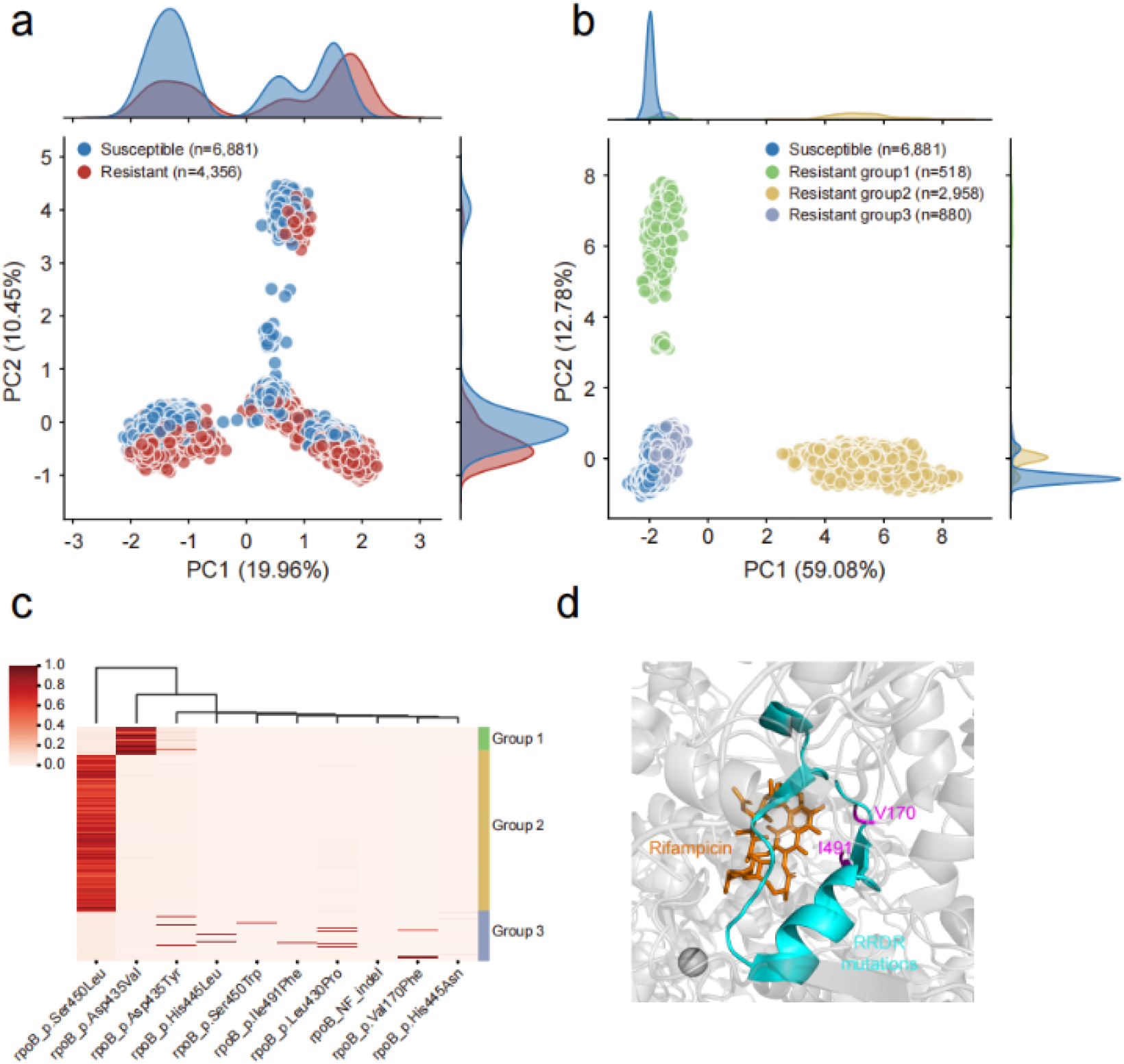
PCA and structural analysis of rifampicin resistance isolates. **a**, PCA of the original features for rifampicin resistance isolates. Marginal density plots are included to illustrate the distribution of the isolates along the first two principal components. **b**, PCA of SHAP values, with resistant isolates color-coded by hierarchical clustering. Marginal density plots are also included, as in panel a. **c**, Heatmap of the top 10 mutations in each resistant group. **d**, Crystal structure of RNA polymerase (PDB: 5UHB) showing the V170F and I491F mutations.

### The framework reflects drug resistance level of isolates

When low-level resistance to first-line drugs occurs, increasing the dosage may help overcome this resistance^27^. We analyzed the relationships between SHAP and MIC values for three first-line drugs using the CRyPTIC dataset (excluding pyrazinamide due to missing data). The isolates were classified into three categories based on MIC value: susceptible, low-level resistant, and high-level resistant following previous studies^26,28,29^. A significant positive correlation between SHAP values of isolates and resistance levels is observed, with the coefficient of determination (R^2^) ranging from 0.44 to 0.69 (Fig.5a-c). Notably, the explainable results from binary classification models appear to contain information related to resistance levels. To further evaluate this, we constructed a ternary classifier using the original features and the SHAP values separately. The classifier trained with SHAP values outperformed the one using original features, showing over 1% improvement (Fig.5d), indicating that SHAP values provide a more effective representation of resistance levels.

**Fig.5.**
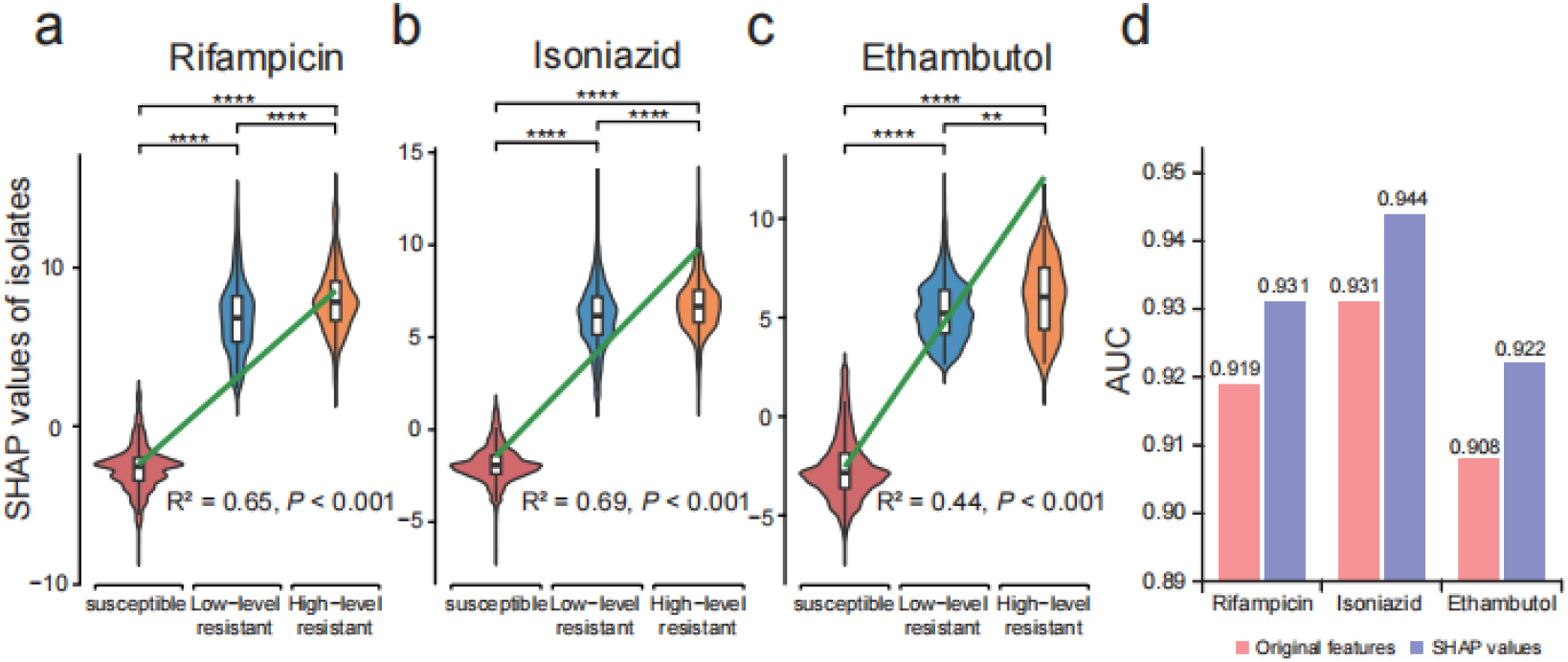
Correlations between the SHAP values and resistance levels of isolates. **a-c**, Violin plots of SHAP values for different resistance levels across the three first-line drugs. Significance was determined using Mann-Whitney test. \*\**P* < 0.01; \*\*\*\**P* < 0.0001. **d**, Performance of the ternary classifier using the original features and SHAP values across different drugs.

## Discussion

xAI-MTBDR predicts MTB drug resistance and identifies resistance-associated mutations at both population and individual levels. To our knowledge, this is the largest study utilizing the MTB WGS dataset (39,145 isolates) to assess the performance and applicability of similar models for drug resistance prediction. The models achieved state-of-the-art performance in predicting resistance to 11 anti-TB drugs, while also identifying new resistance biomarkers, subgrouping drug-resistant isolates, and capturing resistance levels. These insights contribute to a deeper understanding of resistance mechanisms and provide clinical guidance.

Rapid and accurate detection of drug-resistant TB is critical for effective treatment. Our analysis identified potential resistance markers that could expand the current mutation catalogue and provide important clues for further in vivo and in vitro validation to confirm their roles in MTB resistance.

A key strength of xAI-MTBDR is its ability to detect resistance mutations at the individual level, enabling the clustering of isolates with similar resistance mechanisms. For instance, rifampicin-resistant isolates were grouped by distinct mutation patterns, offering a more detailed understanding of resistance. This individual-level insight sets the foundation for more precise prediction models tailored to specific mutation profiles. Although xAI-MTBDR currently focuses on MTB, it is adaptable to other pathogens by substituting resistance-associated genes and reference genomes. This flexibility makes it applicable to a broad range of pathogens, including *Mycobacterium abscessus*^30^, *Staphylococcus aureus*^14^, *Escherichia coli*, and *Klebsiella pneumoniae*^31^, broadening its scope in antimicrobial resistance research.

In summary, we developed an explainable and high-accuracy drug resistance prediction framework designed to facilitate the discovery of new resistance-associated mutations, offering a new perspective on resistance mechanisms at both population and individual levels. As WGS and resistance phenotype data continue to improve, integrating explainable AI methods will enhance prediction accuracy, deepen our understanding of resistance mechanisms, and optimize treatment strategies. This approach has the potential to reduce morbidity and mortality from drug-resistant tuberculosis and other pathogens.

## Methods

### Sequence and DST data

The raw WGS data and corresponding phenotypic DST data for MTB isolates were obtained from three studies: CRyPTIC^18^, Walker et al.^11^, and GenTB^19^. The processing workflow for these datasets is shown in Supplementary Fig.5. The original datasets contained 12,288, 38,223, and 20,379 isolates, respectively. After manual curation to remove unavailable and duplicate isolates, a total of 41,790 isolates were retained. To balance the scale of the datasets, isolates were randomly selected with proportional distributions close to 1:1:1, resulting in 12,285 isolates from CRyPTIC, 14,324 from Walker et al., and 15,181 from GenTB.

Drugs with fewer than 250 isolates in any dataset were excluded^16^. Given the potential overlaps and inconsistencies among the datasets, the DST data were validated and reconciled based on the following principles: (1) consistent DST results from multiple sources were retained; (2) phenotypes from one source were adopted when another lacked data; and (3) inconsistent results were marked as “missing”. The DST categories included susceptible, resistant, and missing. Notably, 12,006 isolates from the CRyPTIC dataset included MIC data, offering a more detailed resistance level beyond the binary resistance phenotype. Table 1 summarizes the proportions of drug-resistant isolates across all datasets.

### Variant calling and feature extraction

All raw WGS data were processed using Clockwork (v0.11.3, https://github.com/iqbal-lab-org/clockwork), a WHO-recommended bioinformatics pipeline for variant calling in Illumina paired-end sequencing data. Quality control filters were applied, retaining only isolates with an average sequencing depth greater than 15, aligned to the H37Rv reference genome (NC000962.3), and with less than 5% of reads mapped to nontuberculous mycobacteria genomes^12^. After filtering, 12,006 isolates from CRyPTIC, 13,750 from Walker et al., and 13,389 from GenTB remained (Supplementary Fig.5). To ensure high-quality mutation data, additional quality control criteria were applied: (1) a minimum sequencing depth of 5x, (2) a fraction of supporting reads of 0.4 or higher, and (3) a genotype confidence percentile (GCP) of 0.5 or greater^18^. Mutation annotation was performed using SnpEff (v5.1d)^32^.

We categorized the mutations identified within 49 genes or promoter regions associated with drug resistance (Supplementary Table 3)^13,14,33,34^ into common (>30 isolates) and rare (≤30 isolates) mutations, following a previous study^20^. While including all rare mutations provides valuable information, their abundance could lead to model overfitting. To address this, rare mutations were classified based on their gene locus (coding, intergenic, and putative promoter regions) and grouped as “derived predictors.” Only common mutations and derived predictors present in at least 30 isolates were retained for the final feature dataset. In total, 788 features were extracted, including 717 common mutations and 71 derived predictors, from 39,145 isolates.

### Model construction and explanation

We combined the strengths of eXtreme Gradient Boosting (XGBoost), random forest, and logistic regression^20,21,35^ to construct a soft voting ensemble model using Python’s Scikit-Learn library (v0.24.2), with voting weights set to [2, 1.5, 1]. The Leave-One-Out (LOO) strategy was applied to train and test the ensemble model for each drug, where one dataset was iteratively selected as the test set while the other two were used for training. Hyperparameters were tuned using RandomizedSearchCV with five-fold cross-validation, followed by manual optimization.

Our framework employs the SHapley Additive exPlanation (SHAP) method, a post-hoc interpretability tool based on game theory, to evaluate each feature’s contribution to the prediction^17^. SHAP is implemented via the *TreeExplainer* for the XGBoost model, which plays the largest role in predicting drug resistance within the ensemble model. SHAP values were calculated for the test dataset of each drug, providing local explanations by assigning SHAP values to each feature for every sample, thus identifying mutations that influence drug resistance at the individual level. Global explanations were derived by averaging the absolute SHAP values of each feature across all samples, highlighting mutations broadly associated with resistance at the population level.

### Identifying potential resistance markers

To accurately identify potential resistance markers, we analyzed the SHAP values of mutations for each correctly predicted isolate, focusing on the most important mutation in each isolate as indicated by the SHAP values from three datasets. Only mutations classified with a final confidence grading of category 1 (associated with resistance) or category 2 (associated with resistance – interim) were considered known resistance-conferring mutations from the WHO mutation catalogue^12,16^. Mutations that are not listed in the WHO mutation catalogue, specifically those not classified in category 1, category 2, category 4 (not associated with resistance – interim), or category 5 (not associated with resistance), but classified in category 3 (uncertain significance) or as unknown mutations, were considered as potential resistance markers. We used Mann-Whitney test to compare differences between the known mutations group and potential mutations group.

### Subgroup the MTB drug-resistant isolates

To investigate the role of SHAP values, we performed principal component analysis (PCA) using the PCA method from sklearn.decomposition for data visualization and compared it with PCA using the original features (where 0 indicates missing mutations and 1 indicates presence). To subgroup the drug-resistant isolates based on SHAP values, we determined the optimal number of clusters using the elbow method (Supplementary Fig.3) and performed clustering using the AgglomerativeClustering method from sklearn.cluster. To identify the key driving mutations within different groups, we applied the Kruskal-Wallis *H* test, and visualized the top 10 statistically significant mutations associated with resistance (*P* < 0.0001) in a heatmap.

### Assessing relationships between SHAP values and resistance levels

To assess the relationship between SHAP values and drug resistance levels, we calculated the sum of SHAP values for each isolate and classified them into three groups: susceptible, low-level resistant, and high-level resistant, based on established criteria^26,28,29^. For rifampicin, MIC ≤ 0.5 μg/ml indicated susceptibility, MIC (0.5, 2] μg/ml indicated low-level resistance, and MIC > 16 μg/ml indicated high-level resistance. For isoniazid, MIC ≤ 0.1 μg/ml indicated susceptibility, MIC (0.1, 3.2] μg/ml indicated low-level resistance, and MIC ≥ 6.4 μg/ml indicated high-level resistance. For ethambutol, MIC < 4 μg/ml indicated susceptibility, MIC [4, 16] μg/ml indicated low-level resistance, and MIC ≥ 32 μg/ml indicated high-level resistance. We used the ggplot2 R package to create violin plots of SHAP values across different resistance levels and employed Spearman analysis to investigate correlations. The Mann-Whitney test was used to compare differences among the three groups.

We developed a ternary XGBoost model (sensitive, low-level resistant, and high-level resistant) using MIC data and trained it separately with the original features and SHAP values from the CRyPTIC dataset. The data was split into training and testing sets with an 8:2 ratio to evaluate model performance.

## Author contributions

Hui Cen was responsible for data collection, analysis, and interpretation, and wrote the manuscript. Peng Zhang assisted with experimental design, particularly in model selection and interpretation, and contributed to the in-depth analysis and interpretation of data. Yunchao Ling participated in result discussions and provided valuable suggestions on data processing and analytical methods. The project was designed by Guoping Zhao. Guoqing Zhang conceived the project, supervised its execution, and edited the manuscript. All authors read and approved the final manuscript.

