## Supplementary figures and tables for "An explainable artificial intelligence framework reveals mutations associated with drug resistance in *Mycobacterium tuberculosis*"

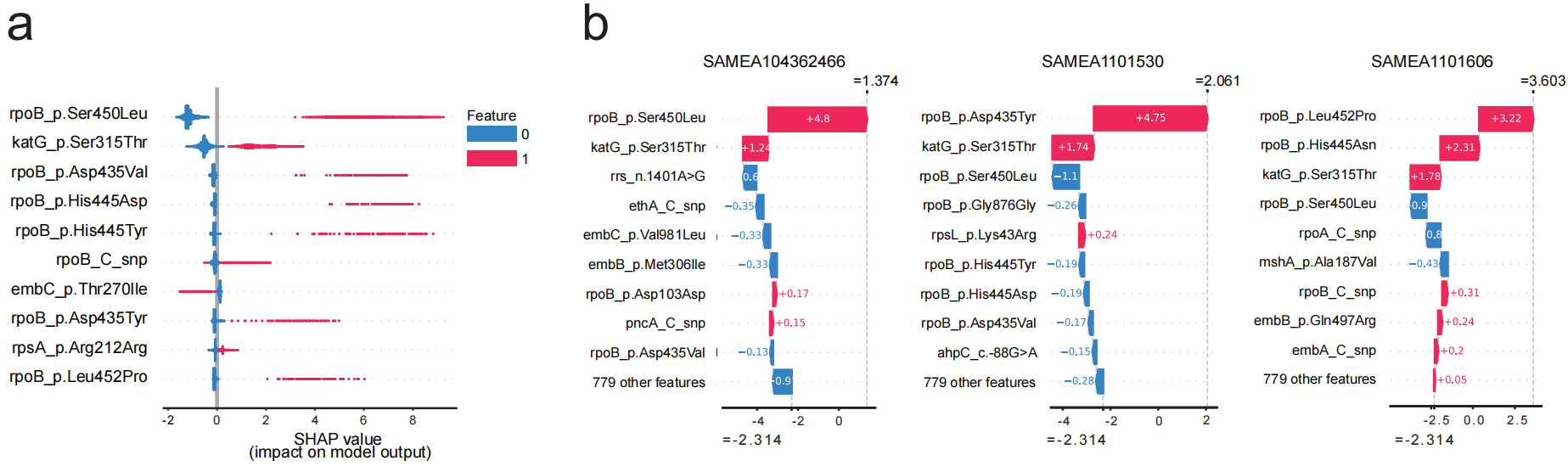

**Figure S1: Population and individual-level explanations in rifampicin isolates.** **a,** Beeswarm plot for population-level explanations. The top 10 mutations are ranked from top to bottom according to their mean absolute SHAP values. The red and blue dots represent isolates with and without this mutation, respectively. The position on the x-axis quantifies the impact of each mutation on the model's prediction. Isolates with certain mutations (red) are mostly on the positive side of the x-axis, indicating that these mutations are associated with higher probabilities of drug resistance. Conversely, isolates missing these mutations (blue) cluster on the negative side of the x-axis, suggesting that their absence is associated with lower probabilities of drug resistance. **b,** Waterfall plots of individual-level explanations. The top 10 mutations are ranked from top to bottom based on their SHAP values. The bottom value of -2.314 represents the base value, which is the average model output over the training data. The sum of the SHAP values for each isolate corresponds to the predicted probability of resistance for that isolate (shown in the upper right corner).

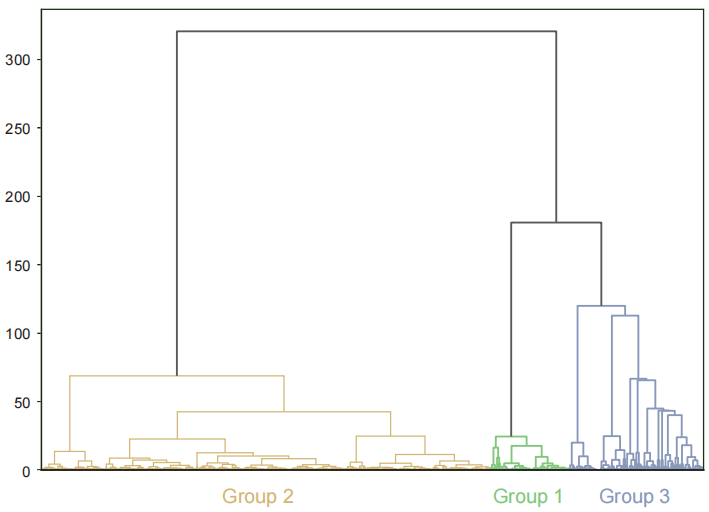

**Figure S2: Dendrogram representing hierarchical clustering of rifampicin-resistant isolates.** The dendrogram illustrates the relationships among different groups of isolates. The vertical axis indicates the dissimilarity or distance between clusters, while the horizontal axis labels the formed groups: Group 1, Group 2, and Group 3. The height at which branches merge reflects the degree of similarity, with shorter branches indicating greater similarity among the groups.

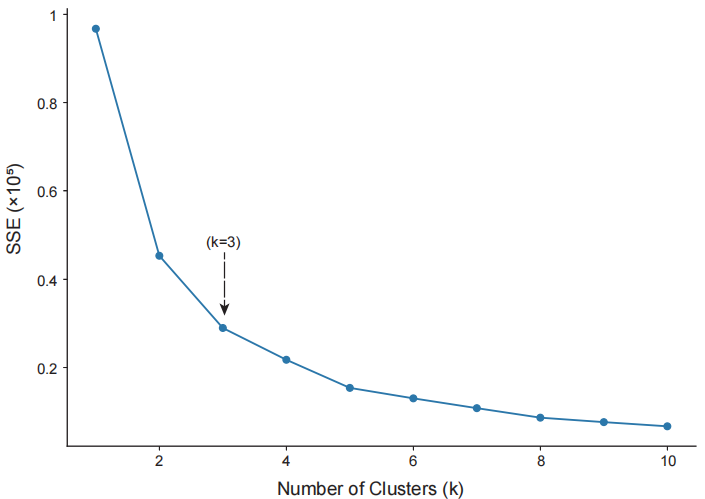

**Figure S3: Elbow plot for determining the optimal number of clusters.** The plot depicts the relationship between the number of clusters k and the sum of squared errors (SSE). The SSE decreases sharply initially but then starts to level off after k=3, forming an "elbow". This suggests that increasing the number of clusters beyond 3 does not significantly improve the partitioning of the data, making k=3 the optimal choice for clustering this dataset.

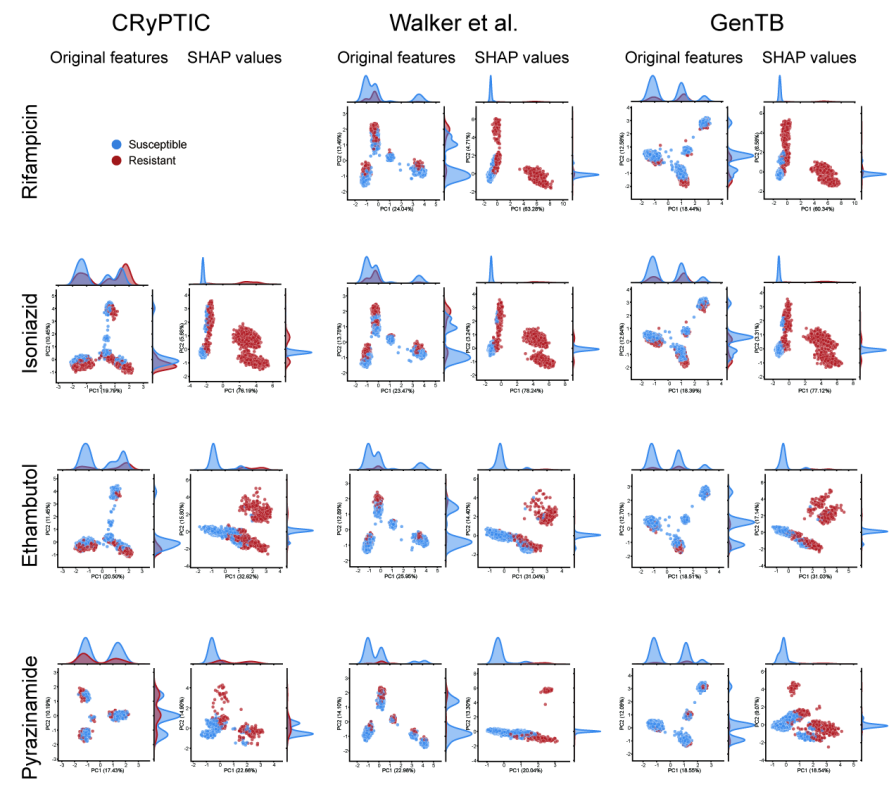

**Figure S4: PCA of original features and SHAP values across different datasets and drugs.**

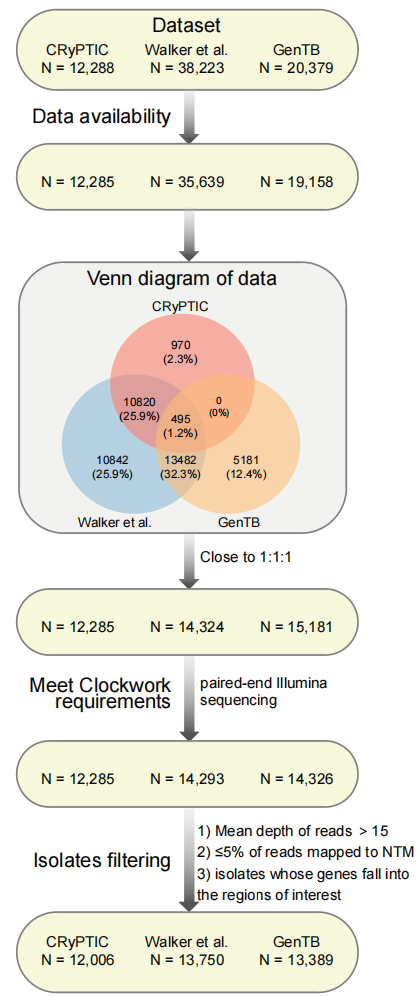

**Figure S5: Workflow of dataset processing.**

**Table S1: 98 important mutations for four first-line drugs identified by individual-level explanations.**

| Drug | Position | Gene | Mutation | WHO Category 1/2 | Potential markers |
| --- | --- | --- | --- | --- | --- |
| rifampicin | 761155 | rpoB | p.Ser450Leu | Yes |  |
| rifampicin | 761110 | rpoB | p.Asp435Val | Yes |  |
| rifampicin | 761139 | rpoB | p.His445Tyr | Yes |  |
| rifampicin | 761140 | rpoB | p.His445Leu | Yes |  |
| rifampicin | 761109 | rpoB | p.Asp435Tyr | Yes |  |
| rifampicin | 761139 | rpoB | p.His445Asn | Yes |  |
| rifampicin | 761140 | rpoB | p.His445Arg | Yes |  |
| rifampicin | 761155 | rpoB | p.Ser450Trp | Yes |  |
| rifampicin | 761139 | rpoB | p.His445Asp | Yes |  |
| rifampicin | 761101 | rpoB | p.Gln432Pro | Yes |  |
| rifampicin | 761095 | rpoB | p.Leu430Pro | Yes |  |
| rifampicin | 761161 | rpoB | p.Leu452Pro | Yes |  |
| rifampicin | 761128 | rpoB | p.Ser441Leu | Yes |  |
| rifampicin | 760314 | rpoB | p.Val170Phe | Yes |  |
| rifampicin | 764817 | rpoC | p.Val483Gly |  | Yes |
| rifampicin | 761110 | rpoB | p.Asp435Gly | Yes |  |
| rifampicin | 761100 | rpoB | p.Gln432Lys | Yes |  |
| rifampicin | 761277 | rpoB | p.Ile491Phe | Yes |  |
| isoniazid | 2155168 | katG | p.Ser315Thr | Yes |  |
| isoniazid | 1674481 | inhA | p.Ser94Ala |  | Yes |
| isoniazid | 1674782 | inhA | p.Ile194Thr |  | Yes |
| isoniazid | 2726141 | ahpC | c.-52C>T |  | Yes |
| isoniazid | 2155168 | katG | p.Ser315Asn | Yes |  |
| isoniazid | 1674263 | inhA | p.Ile21Thr |  | Yes |
| isoniazid | 2726145 | ahpC | c.-48G>A |  | Yes |
| isoniazid | 2155169 | katG | p.Ser315Gly |  | Yes |
| isoniazid | 1674262 | inhA | p.Ile21Val |  | Yes |
| isoniazid | 2726121 | ahpC | c.-72C>T |  | Yes |
| isoniazid | 2726142 | ahpC | c.-51G>A |  | Yes |
| isoniazid | 1674883 | inhA | p.Ile228Val |  | Yes |
| isoniazid | 575909 | mshA | p.Leu188Val |  | Yes |
| ethambutol | 4247429 | embB | p.Met306Val | Yes |  |
| ethambutol | 4247729 | embB | p.Gly406Ser | Yes |  |
| ethambutol | 4247469 | embB | p.Tyr319Ser | Yes |  |
| ethambutol | 4247730 | embB | p.Gly406Asp | Yes |  |
| ethambutol | 4249583 | embB | p.Asp1024Asn |  | Yes |
| ethambutol | 4247730 | embB | p.Gly406Ala | Yes |  |
| ethambutol | 4248002 | embB | p.Gln497Lys | Yes |  |
| ethambutol | 4248003 | embB | p.Gln497Arg | Yes |  |
| ethambutol | 4247431 | embB | p.Met306Ile | Yes |  |
| ethambutol | 4247574 | embB | p.Asp354Ala | Yes |  |
| ethambutol | 4249518 | embB | p.His1002Arg |  | Yes |
| ethambutol | 4247429 | embB | p.Met306Leu | Yes |  |
| ethambutol | 4247431 | embB | p.Met306Ile | Yes |  |
| ethambutol | 4247431 | embB | p.Met306Ile | Yes |  |
| ethambutol | 4247496 | embB | p.Asp328Gly |  | Yes |
| ethambutol | 4247847 | embB | p.Gln445Arg |  | Yes |
| ethambutol | 1416410 | embR | p.Leu313Arg |  | Yes |
| ethambutol | 4247495 | embB | p.Asp328Tyr | Yes |  |
| ethambutol | 4247729 | embB | p.Gly406Cys | Yes |  |
| ethambutol | 4248200 | embB | p.Ile563Leu |  | Yes |
| ethambutol | 4248003 | embB | p.Gln497Pro |  | Yes |
| ethambutol | 4247469 | embB | p.Tyr319Cys | Yes |  |
| ethambutol | 1416699 | embR | p.Leu217Val |  | Yes |
| ethambutol | 4241022 | embC | p.Ala387Val |  | Yes |
| ethambutol | 4247402 | embB | p.Ser297Ala |  | Yes |
| ethambutol | 4247513 | embB | p.Tyr334His |  | Yes |
| ethambutol | 4247728 | embB | p.Glu405Asp |  | Yes |
| pyrazinamide | 2288805 | pncA | p.Ala146Val | Yes |  |
| pyrazinamide | 2289231 | pncA | p.Leu4Ser | Yes |  |
| pyrazinamide | 2289073 | pncA | p.His57Asp | Yes |  |
| pyrazinamide | 2288820 | pncA | p.Gln141Pro | Yes |  |
| pyrazinamide | 2289213 | pncA | p.Gln10Pro | Yes |  |
| pyrazinamide | 2288826 | pncA | p.Val139Ala | Yes |  |
| pyrazinamide | 2289207 | pncA | p.Asp12Ala | Yes |  |
| pyrazinamide | 2289043 | pncA | p.Ser67Pro | Yes |  |
| pyrazinamide | 2289202 | pncA | p.Cys14Gly |  | Yes |
| pyrazinamide | 4039729 | clpC1 | p.Asp326Asn |  | Yes |
| pyrazinamide | 2289213 | pncA | p.Gln10Arg | Yes |  |
| pyrazinamide | 2289040 | pncA | p.Trp68Gly | Yes |  |
| pyrazinamide | 2289072 | pncA | p.His57Arg | Yes |  |
| pyrazinamide | 2288839 | pncA | p.Thr135Pro | Yes |  |
| pyrazinamide | 2288955 | pncA | p.Lys96Thr | Yes |  |
| pyrazinamide | 2289081 | pncA | p.Pro54Leu | Yes |  |
| pyrazinamide | 2288952 | pncA | p.Gly97Asp | Yes |  |
| pyrazinamide | 2289016 | pncA | p.Thr76Pro | Yes |  |
| pyrazinamide | 2289222 | pncA | p.Val7Gly | Yes |  |
| pyrazinamide | 2288727 | pncA | p.Leu172Pro | Yes |  |
| pyrazinamide | 2288844 | pncA | p.Ile133Thr | Yes |  |
| pyrazinamide | 2288790 | pncA | p.Leu151Ser | Yes |  |
| pyrazinamide | 2288937 | pncA | p.Ala102Val |  | Yes |
| pyrazinamide | 2288850 | pncA | c.390_391dupGG |  | Yes |
| pyrazinamide | 2288933 | pncA | p.Tyr103* |  | Yes |
| pyrazinamide | 2289031 | pncA | p.His71Tyr | Yes |  |
| pyrazinamide | 2289090 | pncA | p.His51Arg | Yes |  |
| pyrazinamide | 2289096 | pncA | p.Asp49Gly | Yes |  |
| pyrazinamide | 2289040 | pncA | p.Trp68Arg | Yes |  |
| pyrazinamide | 2289220 | pncA | p.Asp8Asn | Yes |  |
| pyrazinamide | 2289202 | pncA | p.Cys14Arg | Yes |  |
| pyrazinamide | 2289162 | pncA | p.Leu27Pro | Yes |  |
| pyrazinamide | 2288782 | pncA | p.Arg154Gly | Yes |  |
| pyrazinamide | 2289099 | pncA | p.Lys48Thr | Yes |  |
| pyrazinamide | 2288713 | pncA | p.Thr177Pro | Yes |  |
| pyrazinamide | 2288847 | pncA | p.Gly132Ala | Yes |  |
| pyrazinamide | 2289091 | pncA | p.His51Asp | Yes |  |
| pyrazinamide | 2288850 | pncA | c.391dupG |  | Yes |
| pyrazinamide | 2288764 | pncA | p.Thr160Pro | Yes |  |
| pyrazinamide | 2288697 | pncA | p.Leu182Ser | Yes |  |

**Table S2: Distances between drugs and known mutations or potential markers.**

| Drug | Gene | Mutation | Type | Minimum atom distance (Å) |
| --- | --- | --- | --- | --- |
| isoniazid | inhA | p.Ser94Ala | known mutation | 4.0 |
| isoniazid | inhA | p.Ile21Thr | potential marker | 3.8 |
| isoniazid | inhA | p.Ile21Val | potential marker | 3.8 |
| isoniazid | inhA | p.Ile194Thr | potential marker | 3.7 |
| isoniazid | inhA | p.Ile228Val | potential marker | 15.9 |
| ethambutol | embB | p.Leu74Arg | known mutation | 19.9 |
| ethambutol | embB | p.Met306Ile | known mutation | 8.6 |
| ethambutol | embB | p.Met306Val | known mutation | 8.6 |
| ethambutol | embB | p.Met306Leu | known mutation | 8.6 |
| ethambutol | embB | p.Tyr319Ser | known mutation | 9.7 |
| ethambutol | embB | p.Tyr319Cys | known mutation | 9.7 |
| ethambutol | embB | p.Asp328Tyr | known mutation | 10.1 |
| ethambutol | embB | p.Asp354Ala | known mutation | 14.1 |
| ethambutol | embB | p.Gly406Ala | known mutation | 12.6 |
| ethambutol | embB | p.Gly406Asp | known mutation | 12.6 |
| ethambutol | embB | p.Gly406Ser | known mutation | 12.6 |
| ethambutol | embB | p.Gly406Cys | known mutation | 12.6 |
| ethambutol | embB | p.Gln497Arg | known mutation | 16.6 |
| ethambutol | embB | p.Gln497Lys | known mutation | 16.6 |
| ethambutol | embB | p.Ser297Ala | potential marker | 9.1 |
| ethambutol | embB | p.Asp328Gly | potential marker | 10.1 |
| ethambutol | embB | p.Tyr334His | potential marker | 9.1 |
| ethambutol | embB | p.Glu405Asp | potential marker | 9.7 |
| ethambutol | embB | p.Gln445Arg | potential marker | 8.3 |
| ethambutol | embB | p.Gln497Pro | potential marker | 16.6 |
| ethambutol | embB | p.Ile563Leu | potential marker | 43.8 |
| ethambutol | embB | p.His1002Arg | potential marker | 18.0 |
| ethambutol | embB | p.Asp1024Asn | potential marker | 6.1 |

**Table S3: 49 genes or promoter regions associated with drug resistance.**

| Gene/  Promoter | Drug | Description | Type | Start | End | Length |
| --- | --- | --- | --- | --- | --- | --- |
| rrl | amikacin/capreomycin/kanamycin | 23S ribosomal RNA | rRNA | 1473658 | 1476795 | 3138 |
| rrs | amikacin/capreomycin/kanamycin | 16S ribosomal RNA | rRNA | 1471846 | 1473382 | 1537 |
| Rv0678 | bedaquiline/clofazimine | hypothetical protein | protein coding | 778990 | 779487 | 498 |
| tlyA | capreomycin | 16S/23S rRNA (cytidine-2'-O)-methyltransferase TlyA | protein coding | 1917940 | 1918746 | 807 |
| gyrA | ciprofloxacin/ofloxacin/levofloxacin/moxifloxacin | DNA gyrase subunit A | protein coding | 7302 | 9818 | 2517 |
| gyrB | ciprofloxacin/ofloxacin/levofloxacin/moxifloxacin | DNA gyrase subunit B | protein coding | 5240 | 7267 | 2028 |
| ald | cycloserine | L-alanine dehydrogenase | protein coding | 3086820 | 3087935 | 1116 |
| alr | cycloserine | alanine racemase | protein coding | 3840194 | 3841420 | 1227 |
| ddn | delamanid | deazaflavin-dependent nitroreductase | protein coding | 3986844 | 3987299 | 456 |
| fgd1 | delamanid | F420-dependent glucose-6-phosphate dehydrogenase | protein coding | 490783 | 491793 | 1011 |
| fbiA | delamanid | 2-phospho-L-lactate transferase | protein coding | 3640543 | 3641538 | 996 |
| embA | ethambutol | arabinosyltransferase A | protein coding | 4243233 | 4246517 | 3285 |
| embC | ethambutol | arabinosyltransferase C | protein coding | 4239863 | 4243147 | 3285 |
| embR | ethambutol | transcriptional regulator EmbR | protein coding | 1416181 | 1417347 | 1167 |
| mshA | ethionamide | D-inositol 3-phosphate glycosyltransferase | protein coding | 575348 | 576790 | 1443 |
| ethA | ethionamide | monooxygenase EthA | protein coding | 4326004 | 4327473 | 1470 |
| ethR | ethionamide | HTH-type transcriptional repressor EthR | protein coding | 4327549 | 4328199 | 651 |
| ahpC | isoniazid | alkyl hydroperoxide reductase subunit AhpC | protein coding | 2726193 | 2726780 | 588 |
| ahpC promoter | isoniazid | ahpC promoter | promoter | 2726088 | 2726192 | 105 |
| fabG1 | isoniazid | 3-oxoacyl-ACP reductase FabG | protein coding | 1673440 | 1674183 | 744 |
| inhA | isoniazid | NADH-dependent enoyl-[ACP] reductase | protein coding | 1674202 | 1675011 | 810 |
| inhA promoter | isoniazid | inhA promoter | promoter | 1674184 | 1674201 | 18 |
| kasA | isoniazid | 3-oxoacyl-ACP synthase 1 | protein coding | 2518115 | 2519365 | 1251 |
| katG | isoniazid | catalase-peroxidase | protein coding | 2153889 | 2156111 | 2223 |
| ndh | isoniazid | NADH dehydrogenase | protein coding | 2101651 | 2103042 | 1392 |
| oxyR | isoniazid | pseudo | pseudo | 2725571 | 2726087 | 517 |
| embB | isoniazid/ethambutol | arabinosyltransferase B | protein coding | 4246514 | 4249810 | 3297 |
| iniA | isoniazid/ethambutol | isoniazid inductible protein IniA | protein coding | 410838 | 412760 | 1923 |
| iniB | isoniazid/ethambutol | isoniazid inducible protein IniB | protein coding | 409362 | 410801 | 1440 |
| iniC | isoniazid/ethambutol | iIsoniazid inductible protein IniC | protein coding | 412757 | 414238 | 1482 |
| eis | kanamycin | enhanced intracellular survival protein | protein coding | 2714124 | 2715332 | 1209 |
| eis promoter | kanamycin | eis promoter | promoter | 2713785 | 2714123 | 339 |
| rplC | linezolid | 50S ribosomal protein L3 | promoter | 800809 | 801462 | 654 |
| pncA | pyrazinamide | pyrazinamidase/nicotinamidase PncA | protein coding | 2288681 | 2289241 | 561 |
| rpsA | pyrazinamide | 30S ribosomal protein S1 | protein coding | 1833542 | 1834987 | 1446 |
| rpsA promoter | pyrazinamide | rpsA promoter | promoter | 1833380 | 1833541 | 162 |
| panD | pyrazinamide | aspartate 1-decarboxylase | protein coding | 4043862 | 4044281 | 420 |
| clpC1 | pyrazinamide | ATP-dependent protease ATP-binding subunit ClpC | protein coding | 4038158 | 4040704 | 2547 |
| clpP2 | pyrazinamide | ATP-dependent CLP protease proteolytic subunit 2 | protein coding | 2762531 | 2763175 | 645 |
| clpP1 | pyrazinamide | ATP-dependent CLP protease proteolytic subunit 1 | protein coding | 2763172 | 2763774 | 603 |
| folC | para-aminosalicylic acid | folylpolyglutamate synthase FolC | protein coding | 2746135 | 2747598 | 1464 |
| ribD | para-aminosalicylic acid | bifunctional diaminohydroxyphosphoribosylaminopyrimidine deaminase/5-amino-6-(5-phosphoribosylamino)uracil reductase | protein coding | 2986839 | 2987615 | 777 |
| thyA | para-aminosalicylic acid | thymidylate synthase ThyA | protein coding | 3073680 | 3074471 | 792 |
| thyX | para-aminosalicylic acid | thymidylate synthase ThyX | protein coding | 3067193 | 3067945 | 753 |
| rpoB | rifampicin | DNA-directed RNA polymerase subunit beta | protein coding | 759807 | 763325 | 3519 |
| rpoA | rifampicin | DNA-directed RNA polymerase subunit alpha | protein coding | 3877464 | 3878507 | 1044 |
| rpoC | rifampicin | DNA-directed RNA polymerase subunit beta' | protein coding | 763370 | 767320 | 3951 |
| gid | streptomycin | 16S rRNA (guanine(527)-N(7))-methyltransferase RsmG | protein coding | 4407528 | 4408202 | 675 |
| rpsL | streptomycin | 30S ribosomal protein S12 | protein coding | 781560 | 781934 | 375 |
